# Spike initiation properties in the axon support high-fidelity signal transmission

**DOI:** 10.1101/2021.12.13.472435

**Authors:** Mohammad Amin Kamaleddin, Nooshin Abdollahi, Stéphanie Ratté, Steven A Prescott

**Affiliations:** Neurosciences and Mental Health, The Hospital for Sick Children, Toronto, Ontario, Canada; Institute of Biomedical Engineering, University of Toronto, Toronto, Ontario, Canada; Department of Physiology, University of Toronto, Toronto, Ontario, Canada

**Keywords:** axon, excitability, spike initiation, action potential, K_V_1, optophysiology, filter

## Abstract

The axon initial segment (AIS) converts graded depolarization into all-or-none spikes that are transmitted by the axon to downstream neurons. Analog-to-digital transduction and digital signal transmission call for distinct spike initiation properties (filters) and those filters should, therefore, differ between the AIS and distal axon. Here we show that unlike the AIS, which spikes repetitively during sustained depolarization, the axon spikes transiently and only if depolarization reaches threshold before K_V_1 channels activate. Rate of depolarization is critical. This was shown by optogenetically evoking spikes in the distal axon of CA1 pyramidal neurons using different photostimulus waveforms and pharmacological conditions while recording antidromically propagated spikes at the soma, thus circumventing the prohibitive difficulty of patching intact axons. Computational modeling shows that K_V_1 channels in the axon implement a high-pass filter that is matched to the axial current waveform associated with spike propagation, thus maximizing the signal-to-noise ratio to ensure high-fidelity transmission of spike-based signals.

## INTRODUCTION

Nearly all neurons rely on action potentials, or spikes, to transmit information. Variations in the spike initiation process yield diverse spiking patterns^1-3^ associated with distinct coding properties^4-6^. Hodgkin recognized the most fundamental differences and subdivided neurons into three classes: class 1 and 2 neurons spike repetitively during depolarization (consistent with integrators and resonators, respectively) whereas class 3 neurons spike transiently at the onset of depolarization^7^. Those patterns reflect qualitative differences in how voltage-gated currents interact near spike threshold^8, 9^. In class 3 neurons, epitomized by coincidence detectors in the auditory brainstem^10^, a spike occurs only if voltage reaches threshold before subthreshold potassium channels activate or sodium channels inactivate; a high-pass filter is implemented by time- and voltage-dependent reduction of net inward current^11, 12^. Conversely, subthreshold activation of sodium or calcium channels encourages temporal integration and repetitive spiking. Most pyramidal neurons lean toward the integration side of this continuum^13^.

But just as different neurons have evolved spike initiation properties suited to their specific needs, different regions of the same neuron may be specialized to mediate region-specific functions. Spikes are initiated in the axon initial segment (AIS)^14-16^ and transmitted by the axon. During saltatory conduction, where a spike is re-initiated at sequential nodes of Ranvier, the axial current experienced by each node differs from the synaptic input received by the AIS. Processing different signals calls for different filters^17^. Indeed, depolarizing the bleb formed at the cut end of the AIS (<100 µm from soma) or distal axon (>110 µm) produces repetitive and transient spiking, respectively^15^. Other studies have reported transient spiking when stimulating distal axon blebs^18-21^ but damage to the membrane or myelin may affect excitability^22^. Whole-cell recordings from intact distal axons are prohibitively difficult except at boutons or terminals, which may differ in their excitability given their role in synaptic transmission^23^. That said, transient spiking has been consistently observed when injecting current into boutons^24-28^ or the calyx of Held^29-31^. A subset of studies^18, 29, 30^ and older work on peripheral nerves^32, 33^ converted axons from transient to repetitive spiking by blocking potassium currents, linking class 3 excitability to low-threshold K_V_1 or K_V_7 channels^34^. As an interesting side note, squid giant axon also has class 3 excitability^35^, contrary to Hodgkin and Huxley^36^. Kanda et al.^37^ recently succeeded in patching intact nodes of Ranvier in peripheral axons, but they did not study spike initiation and the absence of voltage-gated K channels they reported is hard to reconcile with other findings (see above). Despite growing appreciation for the complexity of axon signaling^38, 39^, axon excitability has remained difficult to study.

We studied axon excitability by evoking spikes optogenetically in the intact distal axon while recording propagated spikes at the soma. Using this non-invasive approach, we confirmed that K_V_1 channels in the axon implement a high-pass filter responsible for transient spiking. This filter matches the axial current associated with spike propagation, thus optimizing transmission of spike-based signals by minimizing the corrupting effects of noise and low-frequency inputs.

## RESULTS

### Spike initiation properties differ qualitatively between the soma/AIS and distal axon

To study axon excitability, we photostimulated the axon of CA1 pyramidal neurons expressing channelrhodopsin-2 (ChR2) while recording from the soma. Perisomatic excitability was tested by photostimulating the soma and by injecting depolarizing current through the recording pipette. Photostimuli were preceded by a photo-prepulse (not shown on most traces) to make photocurrents “square” by removing the inactivating component of ChR2^40^ (**Fig. S1**). Current injection and photostimulation both evoked repetitive spiking when applied to the soma, but photostimulating the axon far (>600 µm) from the soma only evoked a single spike (**Fig. 1A**). Spikes originating near the soma have a smoother onset than spikes propagated from a distant location^21, 41-44^; specifically, spike onset is smooth or abrupt depending on whether the recording site is close (electrically coupled) or far (uncoupled), respectively, from the spike initiation site^45^. As expected, spikes evoked by somatic stimulation had a smooth onset unlike those evoked by axon photostimulation (see gray insets), which had an abrupt onset like spikes evoked by electric field stimulation of the axon (**Fig. 1B**). Though capable of evoking single spikes, electric field stimulation cannot produce the sustained, localized transmembrane current^46^ required to assess spiking pattern. Somatic voltage in our multicompartment model confirmed that smooth-onset spikes originate in the AIS, adjacent to the soma, whereas abrupt-onset spikes originate in the distal axon and propagate to the soma (**Fig. 1C**). Henceforth, we refer to smooth- and abrupt-onset spikes as locally-initiated (LI) and remotely-initiated (RI) relative to the somatic recording site.

**Figure 1.**
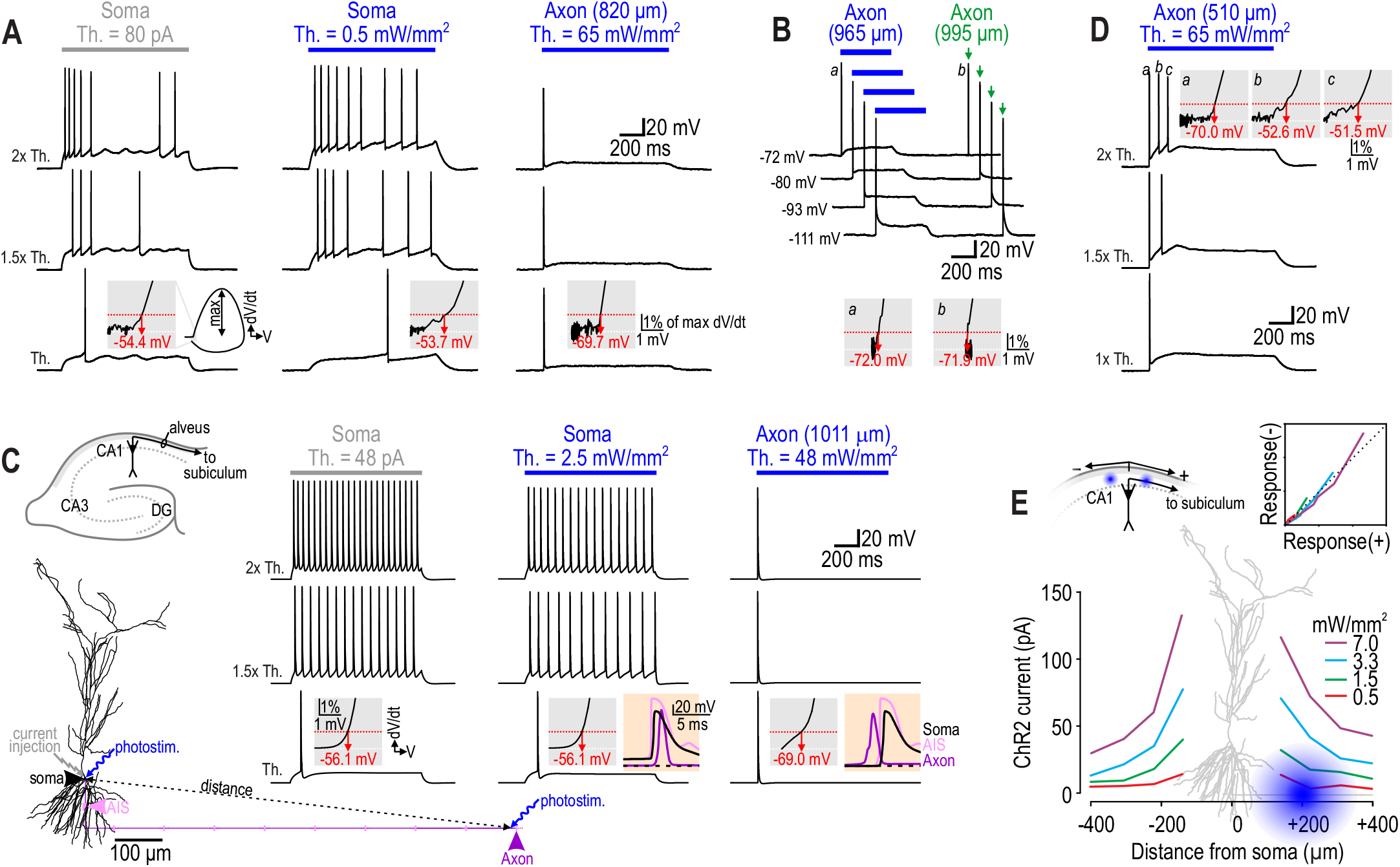
Soma and distal axon respond to sustained depolarization with distinct spiking patterns. **A**. Responses to somatic current injection (*left*) or to photostimulation at the soma (*middle*) or distal axon (*right*) in a CA1 pyramidal neuron. Responses were recorded at the soma. Photostimuli were targeted to a spot ∼20 µm in diameter. Stimulation was repeated at 1, 1.5 and 2ξ threshold (*Th*.) intensity. Gray insets show d*V*/d*t* (normalized to maximum) plotted against *V* to highlight the voltage trajectory at spike onset. Voltage threshold (*red*) is defined as voltage where d*V*/d*t* reaches 1% of its maximum. For somatic stimulation, somatic voltage rises gradually from rest (−70 mV) to threshold (−54 mV); for axon stimulation, somatic voltage abruptly exceeds threshold (−70 mV). **B**. Sample responses from another neuron to axon photostimulation (*blue*) and extracellular electrical pulses (*green*) applied just distal to photostimulation site. Both stimuli evoked a single, abrupt-onset spike that was not eliminated by somatic hyperpolarization. Somatic voltage is indicated beside traces. **C**. Simulations in a multicompartment model. Recording and stimulation sites are indicated on the neuron. Axon is shown in purple (hillock and internodes) and pink (AIS and nodes, enlarged for clarity). Distance between soma and photostimulation site is measured as a straight line (gray; see Methods). Beige insets show enlarged view of voltage at the soma (*black*), AIS (*pink*), and node adjacent to axon photostimulation site (*purple*) plotted against time; dashed line indicates -70 mV. **D**. Sample responses from same neuron as in A, but for axon photostimulation nearer the soma. Proximal axon photostimulation evoked somatic depolarization and multiple spikes, but only the first one had an abrupt onset characteristic of an RI spike (see gray insets). **E**. ChR2 currents evoked by photostimulating the alveus at different distances from the soma toward (+) or away from (−) subiculum are comparable. When plotted relative to each other (*inset on top right*), responses to stimuli in the + or – direction fall along the dotted line representing equivalence even though an axon is present only in the + direction, demonstrating off-target ChR2 activation by light scattered outside the axon target zone. Morphology of our model neuron is superimposed roughly to scale with the *x*-axis. A scattered photostimulus (*blue*) aimed at the alveus (axon) could easily hit nearby dendrites. See Figure S4 for simulations with scattered light.

Axon photostimuli applied closer (<600 µm) to the soma often evoked >1 spike (**Fig. 1D**). This might indicate spatial variation in axon excitability, but only the first spike exhibited an abrupt onset and proximal axon photostimulation also evoked somatic depolarization, which might drive LI spikes. Somatic depolarization could arise via electrotonic spread of current from ChR2 activated in the axon – whose length constant can reportedly reach 800 µm (ref. 47; but see **Fig. S2**) – or because photostimulation targeted to the axon inadvertently activated ChR2 on the dendrites. Consistent with off-target ChR2 activation due to light scattering, responses were insensitive to the precise targeting of photostimuli (**Fig. S3**). To resolve this, we photostimulated the alveus at sites equidistant from the soma toward or away from the subiculum (defined as + and – directions, respectively). Photostimuli in either direction evoked comparable ChR2 currents (**Fig. 1E**) despite absence of the axon in the – direction^48^ due to our slice orientation. All results point to activation of ChR2 outside the intended photostimulation zone. Effects of off-target ChR2 activation were reproduced in simulations when light scattering was added (**Fig. S4**). Without re-tuning the model, light scattering also increased axon photothreshold to values consistent with experiments (see below). All subsequent simulations include light scattering.

These results argue that axons spike exclusively at the onset of depolarization, consistent with class 3 excitability, whereas ability of the AIS to spike repetitively (including at relatively low rates) is consistent with class 1 excitability. But such conclusions hinge on the distinction between LI and RI spikes, which warranted further consideration.

### LI and RI spikes are differentiated by somatic hyperpolarization and local sodium channel blockade

Unlike RI spikes whose initiation in the axon and antidromic propagation to the soma are resilient to somatic hyperpolarization^21, 42^ (see Fig. 1B), LI spikes fail to initiate if local membrane potential is prevented from reaching voltage threshold. Sensitivity to somatic hyperpolarization can, therefore, differentiate LI and RI spikes. As predicted, LI spikes were discouraged by somatic hyperpolarization whereas RI spikes persisted (**Fig. 2A**). Increased somatic leak conductance applied by dynamic clamp had the same effect (**Fig. S5A**). Modest hyperpolarization delayed the onset of LI spikes (see * in Fig. 2A) because the soma’s large capacitance must charge before LI spikes initiate, whereas RI spikes were unaffected except for accentuation of the notch in their rising phase (**Fig. S5B**), which also reflects increased somatic charging time. LI and RI photothresholds were defined as the weakest soma or axon photostimulus able to evoke LI or RI spikes, respectively. All neurons with LI photothreshold measurements (*n*=12) responded with a single LI spike at LI photothreshold (**Fig. 2B**, top trace). When tested at -70 mV, 5 of 6 neurons with RI photothreshold measurements responded to axon photostimuli ≤RI photothreshold with one or more LI spikes (**Fig. 2B**, bottom traces). Hyperpolarizing the soma to -90 mV significantly reduced the number of LI spikes at RI photothreshold (*p* = 0.028; *t*_5_ = 3.05; paired *t*-test) (**Fig. 2B**) and increased LI photothreshold from 0.77 ± 0.12 mW/mm^2^ (mean ± SEM) at -70 mV to 2.53 ± 0.33 mW/mm^2^ at -90 mV (*p* < 0.001; *t*_11_ = - 7.15; paired *t*-test) (**Fig. 2C left**). Somatic hyperpolarization did not affect RI photothreshold (*p* = 0.10; *t*_5_ = 2.47; 83.6 ± 7.0 mW/mm^2^ at -70 mV vs 79.3 ± 5.8 mW/mm^2^ at -90 mV) (**Fig 2C right**). LI spikes evoked by axon photostimulation could conceivably collide with and thus prevent late RI spikes from reaching the soma, but no additional RI spikes were observed when LI spikes were eliminated by hyperpolarization or other means (see below), consistent with occurrence of a single RI spike. At -70 mV, RI photothreshold (83.6 ± 7.0 mW/mm^2^) was a hundredfold greater (*p* < 0.001; Mann-Whitney U test) than LI photothreshold (0.77 ± 0.12 mW/mm^2^), which means scattering just 1% of the light required to evoke an RI spike suffices to evoke LI spikes (see Fig. S4B for discussion).

**Figure 2.**
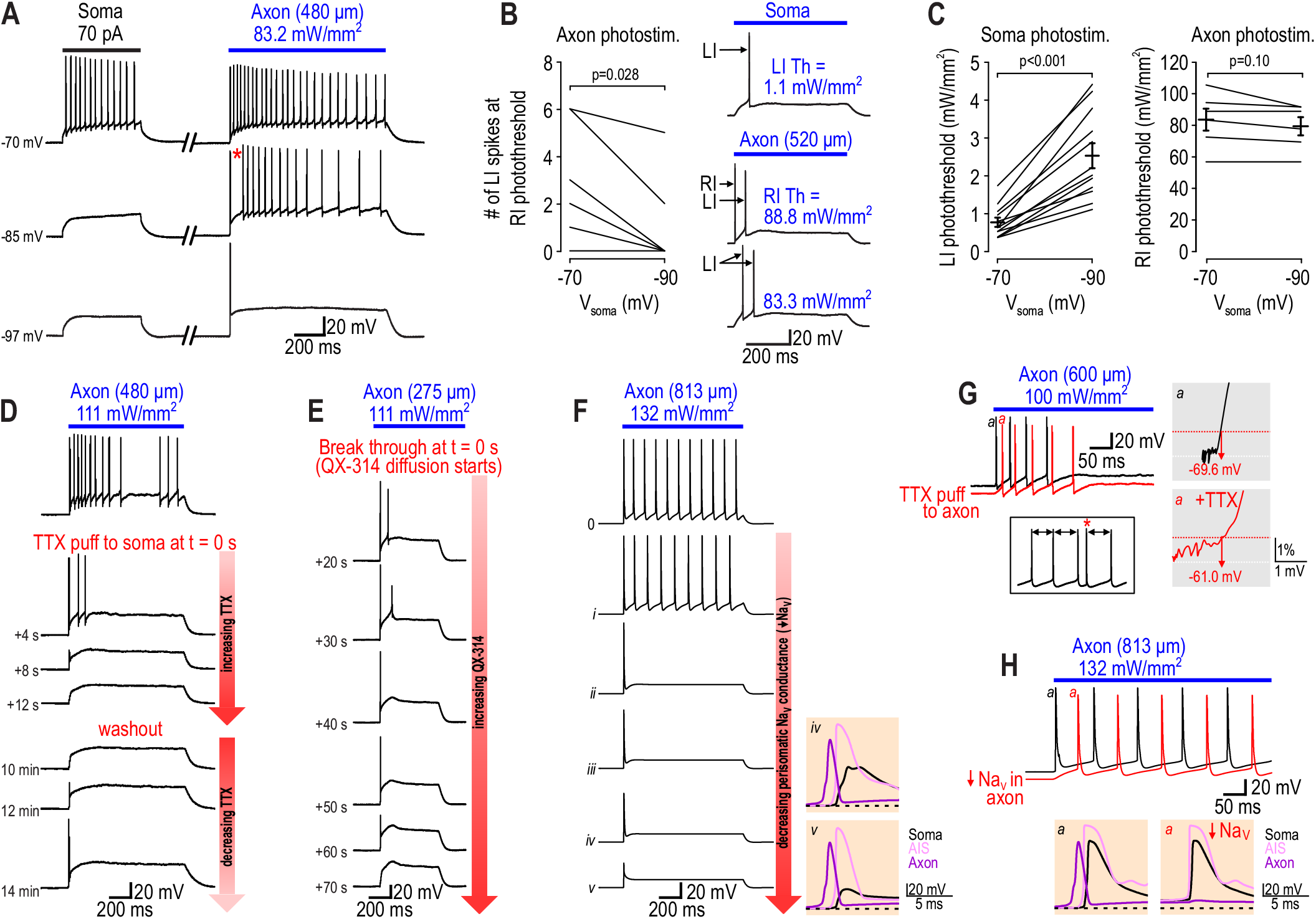
Experimental manipulations confirm perisomatic and axonal origins of LI and RI spikes. **A**. Sample responses with somatic voltage (*V*_soma_) adjusted to values indicated on left. LI spikes were abolished (*bottom*) or delayed *(middle*, *) by somatic hyperpolarization, whereas RI spikes were unaffected. **B**. LI photothreshold was defined as the weakest soma photostimulus able to evoke an LI spike. Only a single LI spike ever occurred at LI photothreshold *(top* trace). RI photothreshold was defined as the weakest axon photostimulus able to evoke an RI spike. LI spikes occurred at intensities ≤RI photothreshold in 5 of 6 neurons tested at -70 mV (*bottom* traces). Hyperpolarization to -90 mV reduced the number of LI spikes (*p* = 0.028; *t*_5_ = 3.05; paired *t*-test), eliminating them in all but two neurons. **C**. Soma and axon photostimulus intensities were titrated to determine LI and RI photothreshold, respectively. Somatic hyperpolarization significantly increased LI photothreshold (*p* < 0.001; *t*_11_ = -7.15; paired *t*-test) from 0.77 ± 0.12 mW/mm^2^ (mean ± SEM) to 2.53 ± 0.33 mW/mm^2^, whereas RI photothreshold was slightly reduced (*p* = 0.10; *t*_5_= 2.00) from 83.6 ± 7.0 mW/mm^2^ to 79.3 ± 5.8 mW/mm^2^. Photostimuli were tested between 360 and 820 µm from the soma (median = 550 µm). **D**. Responses to axon photostimulation were recorded before and at multiple time points after puffing TTX on the soma. Somatic TTX eliminated all LI spikes before eliminating the RI spike in 5 of 10 cells. Unlike LI spikes, the RI spike shrunk gradually and recovered as TTX washed out. The RI spike persisted in 2 cells (see Fig. S5C). **E**.In all cells (*n*=8), infusing QX-314 into the soma eliminated LI spikes before eliminating the RI spike, which shrunk gradually, like in D. **F**. Progressively reducing sodium channel density in our model neuron (conditions *i-v*; see Table S1) caused LI spikes to be lost before the RI spike. Beige insets show voltage at the soma (*black*), AIS (*pink*) and node adjacent to axon photostimulation site (*purple*) to show that the first (RI) spike initiates in the axon but progressively fails to invade the soma as perisomatic sodium channel density is reduced. **G**. In all cells (*n*=3), the RI spike was selectively eliminated by puffing TTX on the axon photostimulation site. Absence of the RI spike allowed the first LI spike to occur earlier, reminiscent of sample traces in Figure 2B. Box shows another example in which an RI spike (*) reset spontaneous LI spikes occurring at regular intervals (arrows). Gray insets show d*V*/d*t* vs *V* plots to illustrate abrupt onset of first (RI) spike before TTX compared with the smooth onset of first (LI) spike after TTX. **H**. Responses to axon photostimulation in the model neuron before and after 90% reduction of axonal sodium channel density. Beige inset shows voltage at soma, AIS and axon to illustrate that if the RI spike does not occur, the first LI spike occurs earlier, consistent with experimental data in G.

We predicted that blocking perisomatic sodium channels would also preferentially eliminate LI spikes. As predicted, Puffing TTX onto the soma promptly eliminated all spikes in most (8 of 10) neurons, but the RI spike was amongst the last eliminated, and was the first to recover as TTX washed out (**Fig. 2D**). In 2 neurons, the RI spike persisted after all LI spikes were eliminated (**Fig. S5C**). In separate recordings, QX-314 applied intracellularly via the patch pipette eliminated all LI spikes before eliminating the RI spike (*n*=8) (**Fig. 2E**). This pattern was reproduced in simulations by progressively reducing sodium channel density in and around the soma (**Fig. 2F)**. In contrast, puffing TTX on the axon photostimulation site selectively eliminated the RI spike (*n*=3) (**Fig. 2G**), allowing the first LI spike to occur earlier. This was reproduced in simulations by reducing sodium channel density in the axon (**Fig. 2H**). These results confirm that LI and RI spikes originate in the AIS and intact axon, respectively.

### K_V_1 channels control axon spiking pattern

Having established that the axon, unlike the AIS, responds to sustained depolarization with a single spike, we proceeded to explore the basis for this spiking pattern. In simulations, reducing K_V_1 density converted the model’s response to axon photostimulation from transient to repetitive spiking (**Fig. 3A**). Blocking K_V_1 channels with low-dose 4-AP converted responses to axon photostimulation to repetitive spiking in all (*n*=5) neurons tested (**Fig. 3B**). Repetitive RI spiking was blocked by applying TTX to the photostimulation site (*n*=2). RI photothreshold was significantly reduced (*t*_7_ = 7.7, *p* < 0.001) from 83.6 ± 7.0 mW/mm^2^ in control conditions (see Fig. 2C) to 3.8 ± 3.1 mW/mm^2^ in 4-AP (*n*=3). Consistent with past observations^42^, spontaneous RI spikes were evident in 4-AP, even during somatic hyperpolarization (**Fig. 3C**). As little as 100 µM 4-AP converted axon photostimulation responses to repetitive spiking in 6 of 8 additional neurons; XE-991 was co-applied in these additional experiments but had no effect on axon excitability when applied by itself. Specifically, when K_V_7 channels were blocked with XE-991, all (*n*=9) neurons continued to spike transiently during axon photostimulation (**Fig. 3D** top) despite obvious changes in their response to somatic current injection (**Fig. 3D** bottom). RI photothreshold was unchanged (*t*_11_ = 0.28, *p* = 0.79, unpaired *t*-test) from 83.6 ± 7.0 mW/mm^2^ in control conditions (see Fig. 2C) to 77.5 ± 19.2 mW/mm^2^ in XE-991 (*n*=7). K_V_7 channels can in principle produce transient spiking (**Fig. S6**) but axon excitability clearly depends on K_V_1 channels in CA1 pyramidal neurons.

**Figure 3.**
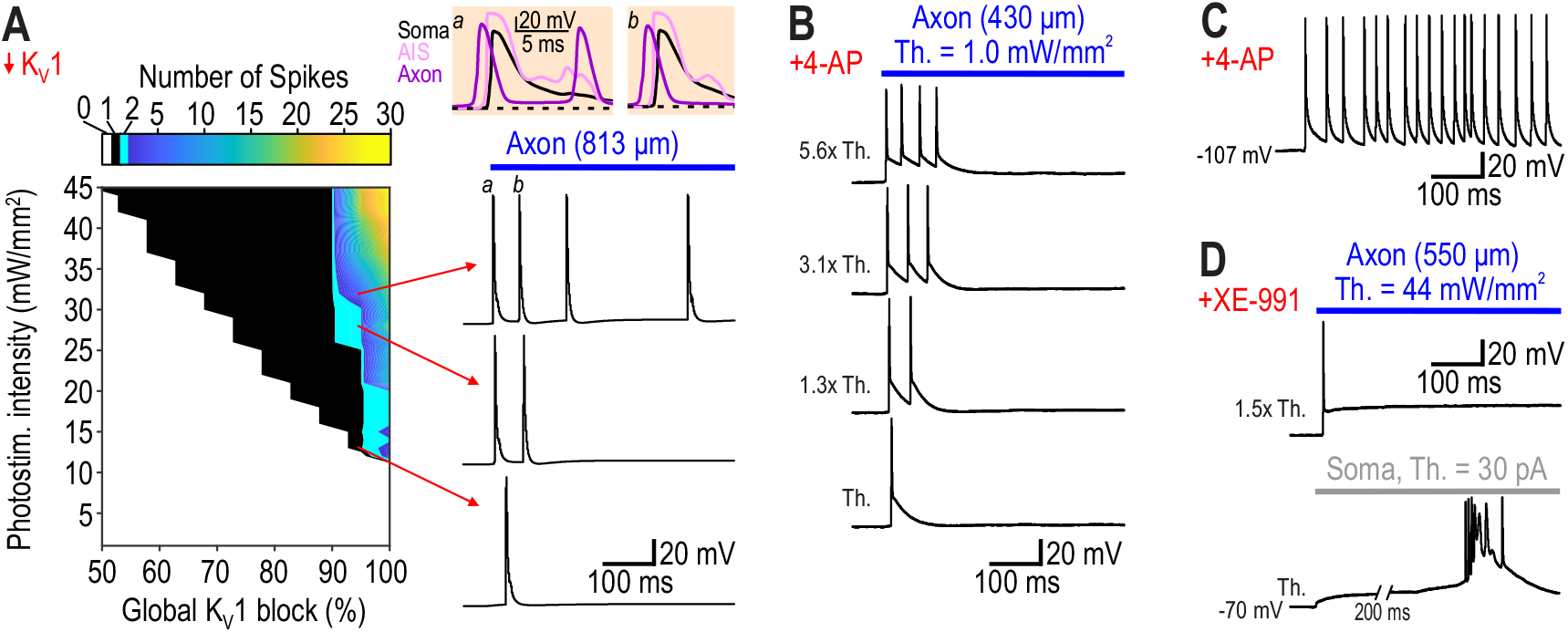
K_V_1 channels dictate spike initiation properties in the axon. **A**. Number of spikes (*color*) during 1 s-long photostimulation of the model neuron depends on photostimulus intensity and K_V_1 density. All spikes recorded at the soma are RI spikes (*beige insets*). The first antidromically conducted RI spike (*a*) triggered a second spike in the AIS that propagated down the axon but not into the soma. This was not observed for subsequent RI spikes (*b*). **B**. All neurons (*n*=5) responded to axon photostimulation with multiple RI spikes after blocking K_V_1 channels with 300 µM 4-AP. The number of RI spikes increased with photostimulation intensity. **C**. Sample of spontaneous RI spiking in 300 µM 4-AP. Spontaneous spiking was not prevented by somatic hyperpolarization. **D**. Blocking K_V_7 channels with 10-30 µM XE-991 never (*n*=9) altered the response to axon photostimulation (*top*) despite clearly affecting responses to somatic current injection (*bottom*).

### Impact of axon spike initiation properties on information transmission

The axon initiates a spike only if its voltage is driven to threshold faster than K_V_1 channels can activate (**Fig. 4A**). Consistent with this explanation, photostimuli applied to the axon as a ramp failed to evoke any spikes (**Fig. 4B**). To compare the input waveform best suited to evoke spikes in the AIS and axon, we injected noisy current (*τ* = 5 ms) into our model neuron in each region and calculated the spike-triggered average (STA) and spike-triggered covariance, and used the latter to plot eigenvectors for the two smallest eigenvalues (**Fig. 4C**). The AIS (left) exhibited a broad monophasic STA and one significant eigenvalue, consistent with a low-pass filter, whereas the axon (right) exhibited a narrow biphasic STA and two significant eigenvalues, consistent with a high-pass filter. Repeating analysis after removing K_V_1 channels confirmed that the high-pass filter is mediated by K_V_1 (**Fig. S7A**). Projecting spike-triggered stimuli onto eigenvectors (bottom panels) shows that spike initiation in the AIS depends exclusively on the amount of depolarization (feature 1), whereas the axon is also sensitive to the rate of depolarization (feature 2). CA1 pyramidal cells behave as integrators because of the low-pass filtering/class 1 excitability of their AIS. The high-pass filtering/class 3 excitability of the electrically segregated axon does not influence neuronal encoding properties but, instead, affects signal transmission.

**Figure 4.**
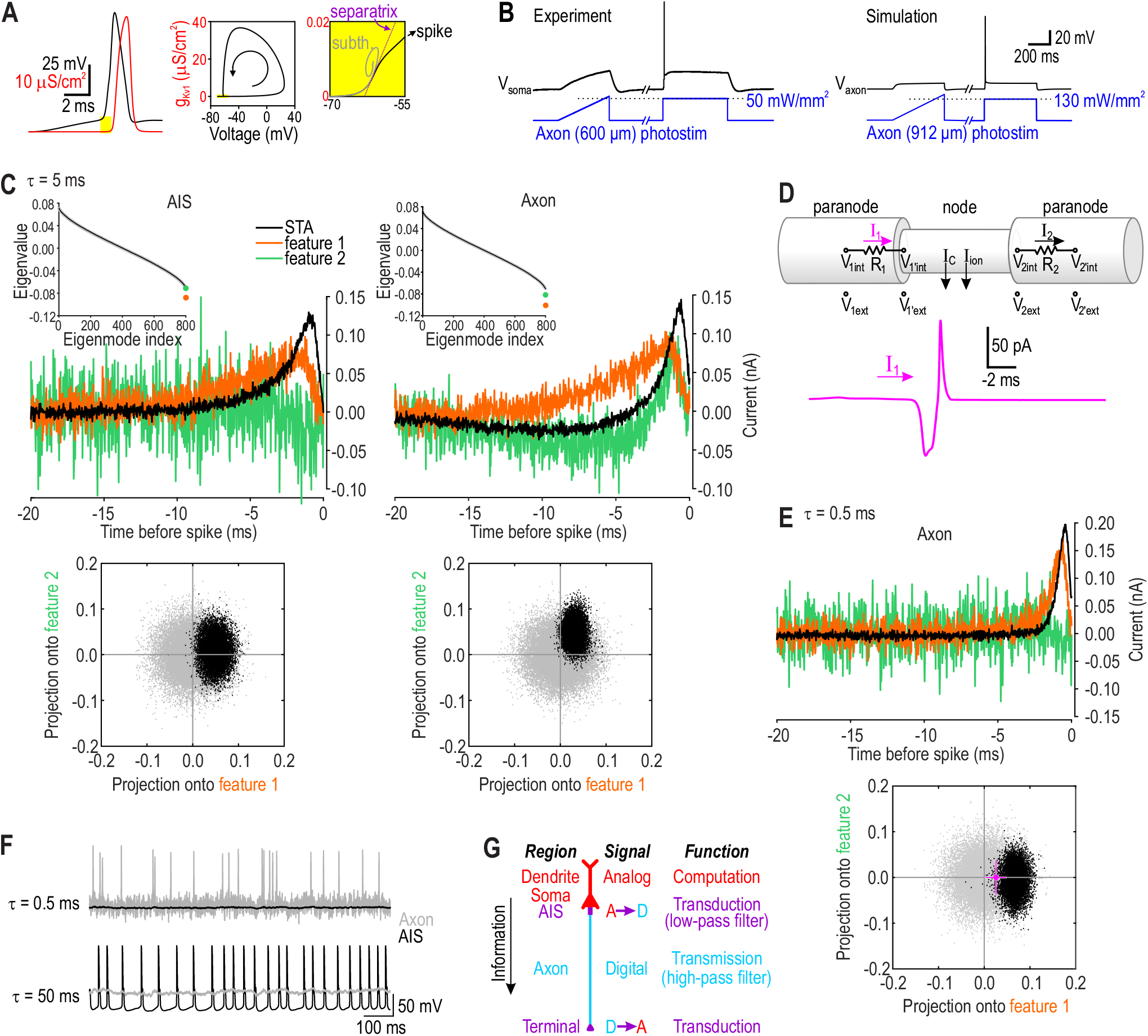
High-pass filtering by K_V_1 channels ensures high-fidelity transmission of spike-based signals. **A**. Axon voltage (*black*) must reach threshold before K_V_1 conductance (*g*_Kv1_, *red*) strongly activates, as evident by plotting voltage against *g*_Kv1_. Zooming in on perithreshold voltages (*yellow*) shows that a spike initiates only if the voltage trajectory (*black*) escapes to the right of the separatrix (*purple*); a just-subthreshold voltage trajectory (*gray*) is shown for comparison. The separatrix reflects a race between fast-activating inward and slower-activating outward currents^9^. **B**. A photostimulus ramp applied to the axon did not evoke a spike in experiments (*left*) or simulations (*right*), despite exceeding RI photothreshold (based on photostimulus steps), demonstrating the importance of rapid depolarization. **C**.Noisy current (−ρ = 5 ms) was injected into the AIS or axon node to calculate the spike-triggered average (STA, black) and spike-triggered covariance. Insets show eigenvalues from spike-triggered stimuli (*circles*) and surrogate data (*black line*). For the AIS (*left*), the STA was broad and monophasic, and there was only one significant eigenvalue (*orange*). For the axon (*right*), the STA was narrower and biphasic, and there were two significant eigenvalues (*orange* and *green*). The latter pattern is characteristic of a high-pass filter. Reducing K_V_1 abolishes the high-pass filter (see **Fig. S7**). Bottom panels show spike-triggered stimuli (*black*) and surrogate data (*gray*) projected onto features (eigenvectors) corresponding to the two smallest eigenvalues. Dimensions of the black cloud relative to the gray cloud reflect which stimulus features influence spike initiation, and argue that spike initiation in the AIS depends solely on the amount of depolarization whereas spike initiation in the axon depends on the amount and rate of depolarization. **D**. Pink trace shows axial current experienced by a node when a spike occurs at the immediate upstream node. The time axis is reversed to correspond to left-to-right flow of axial current in schematic (*top*), which illustrates how axial current is calculated (see Methods for details). **E**. Analysis in C was repeated for the axon using faster noise (1 = 0.5 ms). STA was narrower, consistent with axial current waveform in D. There was only one significant eigenvalue, suggesting that rate of depolarization is unimportant, but this is because all stimulus fluctuations are fast; indeed, these fluctuations are too fast to evoke spikes in the AIS (see F). Intermediate noise (1 = 5 ms) offers some fluctuations fast enough to evoke spikes in the axon and other, slower fluctuations able to evoke spikes in the AIS. The axial current waveform (from D) projected onto the feature space (*pink*, mean ± standard deviation for 10 trials) differs significantly from 0 (*t*_9_=4.56, *p*=0.0014, one-sample *t*-test). **F**. Sample responses to fast (1 = 0.5 ms) and slow (1 = 50 ms) noise. For fast noise (*top*), the axon spiked vigorously and exhibited large voltage fluctuations between spikes, whereas the AIS was quiescent and its voltage fluctuations were blunted. For slow noise (*bottom*), the axon was quiescent whereas the AIS spiked regularly despite input fluctuations. Since input to the AIS normally arrives via the soma and axon hillock, noise was injected into the soma to assess the AIS response. Noise was injected into the axon to assess the axon response. χρ = 0.22 nA and µ = 0.1 nA for all cases except slow noise applied to the soma, where µ was reduced to 0.05 nA to reduce the spike rate. **G**. Cartoon depicting information flow through a neuron. The AIS transduces graded depolarization into spikes using a low-pass filter, but those spikes are transmitted by the axon using a high-pass filter.

To explore signal transmission, we used our model to measure the input experienced by a node during spike propagation – the input each node must respond to by initiating its own spike. Unlike the slow, sustained synaptic depolarization received by the AIS (**Fig. S7B**), nodes experience short but intense pulses of axial current (**Fig. 4D**). That pulse is faster than the axon STA in Figure 4C, but the STA also depends on the autocorrelation time of the noise. As predicted, testing the axon with faster noise (*τ* = 0.5 ms) revealed a faster STA (**Fig. 4E**) that matches the axial current kinetics. When stimulated with fast noise (**Fig. 4F**, top), low-pass filtering due to the soma’s large capacitance prevented any spikes from initiating at the AIS (gray), unlike the axon’s vigorous response (black). Slow noise (*τ* = 50 ms) produced the opposite pattern (**Fig. 4F**, bottom), confirming that the axon and AIS are tuned very differently, consistent with the inputs they are required to process.

## DISCUSSION

Axonal excitability was studied by optogenetically evoking spikes in the intact axon of CA1 pyramidal neurons. All spikes were recorded at the soma but their origin at/near the photostimulation site was verified in multiple ways. This non-invasive approach confirmed that intact axons spike transiently during sustained depolarization, consistent with class 3 excitability. We show further that axons spike transiently because of K_V_1 channels and that this K_V_1-mediated high-pass filter has important consequences for digital signal transmission (**Fig. 4G**).

Axons reliably propagate spikes^49-51^ but the biophysical basis for that reliability (*e*.*g*. high sodium channel density^19^) is hard to reconcile with the rarity of spike originating ectopically in the axon. Matching the nodal spike initiation filter to the axial current waveform associated with an incoming spike (originating at the AIS) optimizes spike propagation by minimizing the chance of noise and other (slower, weaker) inputs evoking spikes. Interestingly, K_V_1 channels reduce the energy efficiency of axon spikes^52^ but their ubiquity in axons suggest that they serve an important function worthy of the metabolic cost. Analog signaling in axons – modulation of spike-triggered synaptic output by simultaneously transmitted subthreshold depolarization^53-55^ – does not diminish the importance of digital signaling but, rather, presents additional challenges that axons have evidently solved. Calcium influx and transmitter release depend on spike width^56, 57^, which itself depends on subthreshold voltage^54, 55, 58^, but this amounts to modulating digital-to-analog transduction rather than true analog signaling (like graded transmission at ribbon synapses^59^). Slow, subthreshold voltage signals must pass along the axon without corrupting the spike trains traveling along the same axon. By preventing slow signals from evoking spikes (see Fig. 4F), K_V_1 channels support the frequency-division multiplexing required for this hybrid signaling. Spike width can also be modulated neurochemically without new spikes being added^60^. In other words, the attractor state representing a spike can vary (manifesting changes in spike width) while the presence/absence of a spike remains a binary distinction. The high-pass filter maintains that distinction by keeping attractor states – quiescence and spike – broadly separated.

Re-initiation of spikes at each node of Ranvier is typically viewed as “boosting” so that the signal can travel farther than in a passive cable^61^. Boosting also occurs in dendrites, where voltage-gated channels help analog signals reach the soma^62^. But unlike in dendrites, saltatory conduction in myelinated axons also constitutes a signal restoration process wherein axial current (analog) in the internode is converted to a spike (digital) in the node, which in turn produces axial current (analog). This A/D/A motif alleviates noise accumulation^63^. Matching the nodal spike initiation filter to the axial current waveform associated with spike propagation helps maximize the signal-to-noise ratio^17^. That the AIS employs a different filter to process the slow-fluctuating synaptic depolarization it receives is simply good design.

Digital signals are well suited for transmitting information reliably but analog signals are better for efficient computing^63, 64^. Neurons exploit both, conducting analog computations in their dendrites, soma and AIS, and then transmitting signals in a digital format. Indeed, spikes cost more energy and convey less information than graded potentials^65^, but these penalties are offset by the benefits of reliable transmission. Analog-to-digital transduction at the AIS dictates neuronal spiking patterns and coding properties; plasticity at the AIS has, accordingly, attracted increasing attention^66-68^. Converting spike trains into synaptic transmitter release at the other end of the axon offers a similarly important locus for plasticity. Transmitting spikes along the intervening axon may seem mundane by comparison, but doing so with high fidelity despite noisy conditions, noisy components, and other biological constraints^69^ should not be taken for granted. Nonlinear interaction between sodium and K_V_1 channels enables coincidence detection when that competition occurs in the AIS, but the same competition occurring in the axon ensures high-fidelity transmission of digital signals. Proper neurological function hinges on this subtle yet elegant aspect of axonal design.

## MATERIALS AND METHODS

### Experiments

#### Animals and slice preparation

All procedures were approved by the Animal Care Committee at The Hospital for Sick Children. To express ChR2 in CA1 pyramidal neurons, CamKII (Ca^2+^/Calmodulin-dependent protein kinase IIα)-Cre mice (JAX# 005359) were crossed with floxed ChR2-eYFP (Ai32) mice (JAX# 012569). Adult mice of either sex aged 6-8 weeks were deeply anesthetized with isoflurane and decapitated. The brain was rapidly removed and placed in the ice-cold oxygenated (95% O_2_ and 5% CO_2_) sucrose-substituted artificial cerebrospinal fluid (aCSF) containing (in mM) 252 sucrose, 2.5 KCl, 2 CaCl_2_, 2 MgCl_2_, 10 D-glucose, 26 NaHCO_3,_ 1.25 NaH_2_PO_4,_ and 5 kynurenic acid. Parasagittal brain slices (300-400 µm) were prepared using a VT-1000S microtome (Leica) and kept in oxygenated sucrose-substituted aCSF for 30 minutes at room temperature before being transferred to normal oxygenated aCSF (126 mM NaCl instead of sucrose and without kynurenic acid) until recording. Slices were transferred to a recording chamber constantly perfused at ∼2 ml/min rate with oxygenated (95% O_2_ and 5% CO_2_) aCSF heated to 31±1 °C using an automatic temperature controller (Warner Instruments) and viewed with an Examiner.A1 microscope (Zeiss) using a 40x, 0.75 NA water immersion objective (N-Achroplan, Zeiss) and IR-1000 Infrared CCD camera (Dage-MTI). Cells were visualized with a long-pass filter (OG590) in the transmitted light path to avoid activating ChR2 while patching.

#### Recording

Pyramidal neurons in the CA1 region of hippocampus were recorded in the whole-cell configuration with >75% series resistance compensation using an Axopatch 200B amplifier (Molecular Devices). Glass pipettes with 5-7 MΩ resistance were pulled from borosilicate fire polished glass capillaries (O.D.: 1.5 mm, I.D.: 0.86 mm; Sutter Instrument) prepared using a PC-10 dual-stage micropipette puller (Narishige). The resting membrane potential, cell capacitance, and input resistance of the neurons were continuously monitored during the experiments. Neurons that experienced a >20% change in either criterion were excluded from analysis. The patch pipette solution contained (in mM) 125 KMeSO_4_, 5 KCl, 10 HEPES, 2 MgCl_2_, 4 Adenosine triphosphate (ATP; Sigma-Aldrich), and 0.4 mM Guanosine triphosphate (GTP; Sigma-Aldrich); pH was adjusted to 7.2-7.3 with KOH. Reported values of membrane potentials are corrected for a liquid junction potential of 9 mV. Neuronal responses were low-pass filtered at 2 kHz and digitized at 20 kHz using a Power1401 computer interface and Signal 6.02 software (Cambridge Electronic Design). A virtual shunt was applied using the dynamic clamp capabilities of Signal 6.02; see Figure S5 for parameter values.

#### Stimulation

Current was injected into the soma as square steps via the patch pipette. The minimum current required to evoke a spike (i.e. threshold, *Th*) is indicated on figures. For optogenetic stimulation, a 455 nm LED was used to deliver blue light through the epifluorescent port of the microscope. Photostimulation was limited to a spot ∼20 µm in diameter by partially closing the field diaphragm in the epifluorescent light path. Light intensity at the specimen plane was measured with a S170C photodiode power sensor and PM100D power meter (ThorLabs). Photostimulus intensity was controlled jointly through the Dual OptoLED controller (Cairn Research) and by adding neutral density filters to achieve smaller increments at low intensities. Photostimulus steps were preceded by a strong 200 ms-long pre-pulse to inactivate ChR2 prior to the step^40^ (see Fig. S1). Extracellular electrical stimulation was applied using a DS3 Isolated Current Stimulator (Digitimer) to deliver constant current via bipolar electrodes positioned across the axon bundle (alveus) just distal to the photostimulation site. Pulses 100 μs in duration were increased in amplitude (0.5-3.0 mA) until spikes were recorded. Timing of all stimuli was controlled by Signal 6.02.

The distance to axon stimulation sites was reported as the shortest distance (i.e. straight line) between the photostimulation site and the soma because the axon was not visualized and its path could not, therefore, be traced. The same approach was used in the model neuron for consistency with experiments. In simulations, the length along the axon 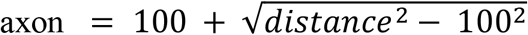, which means distance underestimates length along the axon between 10 and 25% for distances of 1000 to 350 µm, respectively. The underestimation in experiments is even greater due to axon tortuosity and changes in depth that are unaccounted for.

#### Drug application

In all experiments, fast synaptic transmission was blocked by bath application of 10-20 μM 6-Cyano-7-nitroquinoxaline-2,3-dione disodium salt (CNQX; Hello Bio), 40 μM D-2-amino-5-phosphonovalerate (APV; Hello Bio), and 10 μM bicuculline methiodide (Hello Bio). Control experiments confirmed that no component of photoevoked responses was synaptically mediated. To block voltage-gated Na^+^ channels, 10-20 μM tetrodotoxin citrate (TTX; Alomone Labs) dissolved in HEPES-buffered aCSF was applied for 20-1000 ms using a puff pipette positioned above the region of interest. Puff pressure was adjusted to 2-4 psi using a picospritzer (Toohey Company). To block voltage-gated Na^+^ channels intracellularly, 420 μM N-(2,6 Dimethylphenylcarbamoylmethyl) triethylammonium chloride (QX-314; Tocris) was added to the intracellular solution in the patch pipette. The pipette tip was always filled with a drug-free solution. To block voltage-gated K^+^ channels, 100-300 μM 4-aminopyridine (4-AP, Sigma-Aldrich) and 10-30 μM XE-991 dihydrochloride (Abcam) were added to the bath. At concentrations ≤300 μM, 4-AP selectively blocks K_V_1.1, 1.2, 1.5 and 2.1 (ref. 70). We bath applied 4-AP and XE-991 because, consistent with past observations^71^, preliminary testing suggested that blockade of K_V_ channels occurred slowly and, therefore, that puffing the drugs was ineffective.

### Statistical analysis

Kolmogorov-Smirnov tests were used to confirm that distributions were Gaussian. All tests were conducted with SigmaPlot 11 (SYSTAT). Details are reported in the text and figure legends.

### Modeling

A multicompartment model of a mouse CA1 pyramidal neuron with a detailed myelinated axon was constructed using NEURON v8.0 (ref. 72). This model is based on the CA1 pyramidal neuron model described by Shah et al.^73^, but changes were made to its morphology and ion channel densities. All model code will be uploaded to ModelDB and prescottlab.ca upon publication.

#### Morphology

The cell dimensions were uniformly shrunk to account for the difference in the average size of rat (original model) and mouse (experiments) CA1 pyramidal neurons^74^. The reconstructed model reproduces morphological features (e.g. total surface area, total dendritic length, somatic surface area) of mouse CA1 pyramidal neurons^75^ (see **Table 1**). The axon used by Shah et al.^73^ was replaced by a more detailed axon containing an axon hillock, AIS, myelin sheaths, and nodes of Ranvier. Inside the myelin sheath, each internode section consisted of the paranode, juxtaparanode and internode regions^76^ (see **Table 2**). The axon had 17 nodes and a total length of ∼1800 µm.

**Table 1.** Morphological properties of the model neuron compared with published values.

|  | Values for our reconstructed model | Values reported previously | Reference |
| --- | --- | --- | --- |
| Cell body area ($\mu\text{m}^2$) | 115.9 | 137±21 | 75 |
| Total dendritic length ($\mu\text{m}$) | 8275 | 8000 | 74 |
| Main apical shaft diameter ($\mu\text{m}$) | 1.69 | 1.82±0.35 | 75 |
| Apical dendrite branch diameter ($\mu\text{m}$) | $\sim 0.8$ | 0.8±0.14 | 75 |
| Apical dendrite terminal diameter ($\mu\text{m}$) | $\sim 0.6$ | 0.6±0.28 | 75 |
| Number of primary basal dendrites | 4 | 3.04±1.12 | 75 |
| Primary basal dendrite diameter ($\mu\text{m}$) | $\sim 1.3$ | 1.19±0.28 | 75 |
| Higher-order basal dendrite branch diameter ($\mu\text{m}$) | $\sim 0.6$ | 0.6 | 75 |
| Axon hillock diameter ($\mu\text{m}$) | 0.834 | 1.18±0.35 | 75, 85 |
| Total surface area ($\mu\text{m}^2$) | 19631 | 18000 | 74 |

**Table 2.** Morphological parameters of the model’s axon. PNJ, Paranodal junction; JXP, juxtaparanode; IND, internode; AIS, axon initial segment.

| Parameter | Value ($\mu\text{m}$) | Reference |
| --- | --- | --- |
| AIS length | 15 | 77, 86 |
| AIS diameter | 0.334 | 86 |
| Outer diameter of the axon | 0.65 | 87 |
| Internode length | 100 | 76, 77, 86 |
| Total axon length | 1800 |  |
| Node diameter | 0.234 | 76, 86 |
| Node length | 1 | 76, 77, 86 |
| PNJ diameter | 0.5 | 88 |
| PNJ length | 3 | 76, 86 |
| JXP diameter | 0.5 | 88 |
| JXP length | 5 | 76, 86 |
| IND diameter | 0.5 | 88 |
| IND length | 83 | 76, 86 |
| PNJ peri-axonal space width | 0.0018 | 76 |
| JXP peri-axonal space width | 0.008 | 76 |
| IND peri-axonal space width | 0.008 | 76 |
| Number of myelin lamella | 15 | 76 |

#### Passive properties

The specific membrane capacitance was 1 µF/cm^2^ except in myelinated regions, where it was 0.1 µF/cm^2^. For all simulations, the resting membrane potential and temperature was set to the same value as in experiments, 31°C. The axial resistivity (*R*_i_) was 150 Ω.cm for soma, axon hillock and dendrites and 50 Ω.cm for the AIS and axon. Membrane leak conductance was 0.028 mS/cm^2^ for soma and dendrites and 1 mS/cm^2^ for nodes (see **Table 3**).

**Table 3.** Passive electrical properties of model neuron. PNJ, Paranodal junction; JXP, juxtaparanode; IND, internode; AIS, axon initial segment. Adjustments to reproduce experimental data are indicated by “→”.

| Parameter | Value | Reference |
| --- | --- | --- |
| Soma/dendrite/axon hillock/AIS capacitance | 1 $\mu\text{F}/\text{cm}^2$ | 77, 89 |
| Node capacitance | 1 $\mu\text{F}/\text{cm}^2$ | 76, 77 |
| Myelin capacitance per lamella membrane | 0.1 $\mu\text{F}/\text{cm}^2$ | 76, 90 |
| Axon/AIS axial resistivity | 50 $\Omega\cdot\text{cm}$ | 73, 77, 89 |
| Soma/dendrites/axon hillock axial resistivity | 150 $\Omega\cdot\text{cm}$ | 73, 89 |
| Soma/dendrites/axon hillock/AIS specific membrane (leak) conductance | 0.035 → 0.028 $\text{mS}/\text{cm}^2$ | 73, 79 |
| Node membrane (leak) conductance | 1 $\text{mS}/\text{cm}^2$ | 77 |
| Myelin membrane (leak) conductance per lamella membrane | 1 $\text{mS}/\text{cm}^2$ | 76, 90 |
| PNJ membrane (leak) conductance | 1* → 0.61* $\text{mS}/\text{cm}^2$ | 76, 90 |
| JXP/IND membrane (leak) conductance | 0.1* → 0.087* $\text{mS}/\text{cm}^2$ | 76, 90 |
| Leak reversal potential | -70 mV | 73 |

\*scaled by axon diameter/fiber diameter = 0.77

#### Active properties

Conductances included in our model were Na_V_1.6, K_V_7, K_V_1, delayed rectifier K^+^, A-type K^+^, AHP, H current, and T-type Ca^2+^ current. Channel kinetics were taken from published models (see **Table 4** for ModelDB accession numbers). The Na^+^ channel kinetics differ between the soma and axon to include slow inactivation. The A-type K^+^ channel in the distal and proximal apical dendrites had different activation properties: vhalfn = -1 mV or 11 mV for distal and proximal dendrites, respectively. Guided by previously described spatial variations in conductance densities, the density of each conductance for each region was adjusted to reproduce our experimental results (see **Table 4**). In brief, the density of Na_V_1.6 in the AIS and nodes was 13x and 3x higher, respectively, than in soma^77^. Ion channel densities were generally low in internodes. Density of K_V_7 channels was higher in nodes compared to internodes, and the density of K_V_1 channels was higher in the juxtaparanode^78^. The model included ChR2 modeled as described by Foutz et al.^79^ (see below).

**Table 4.**
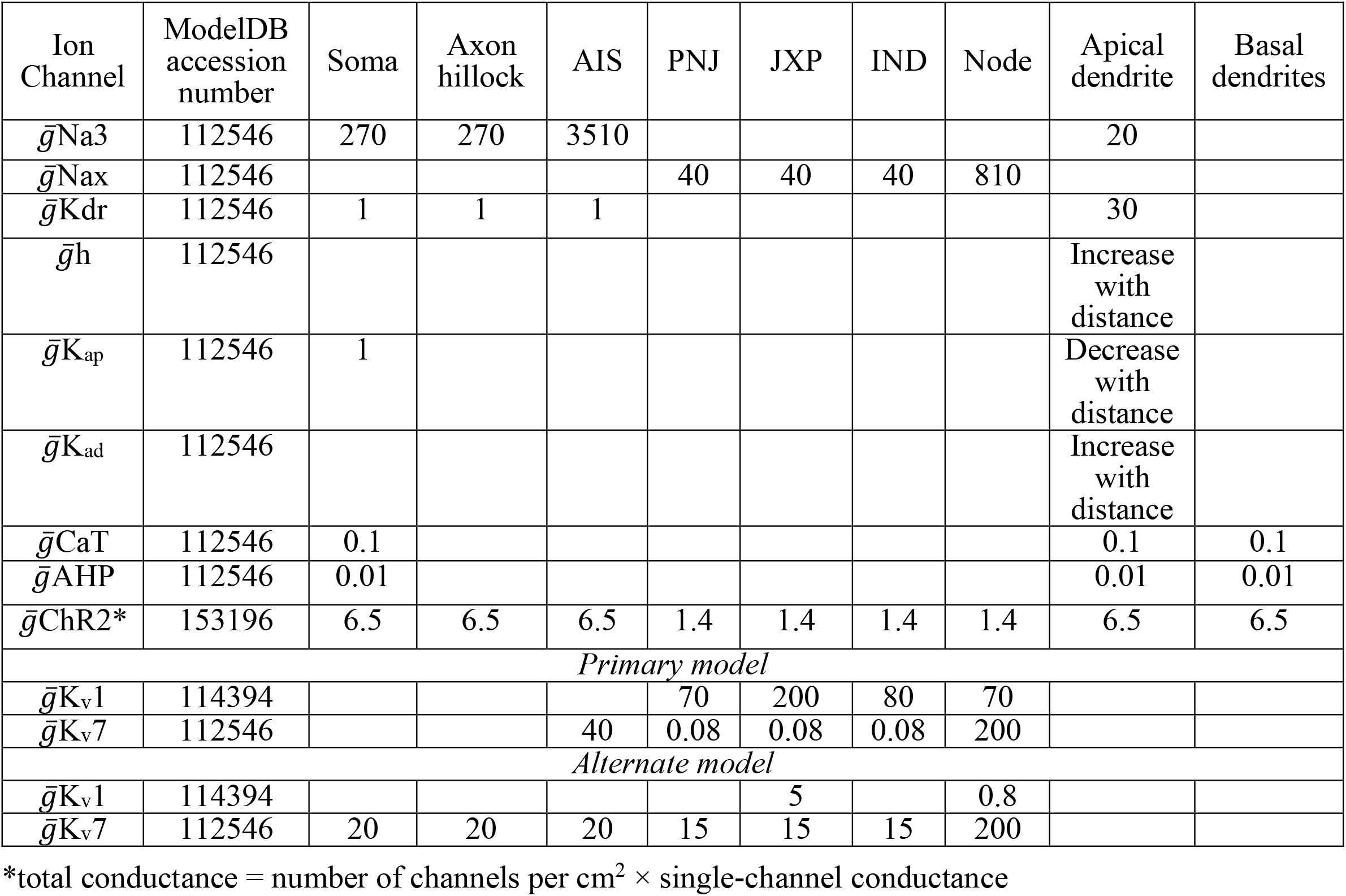
Type, ModelDB accession number, and density (in mS/cm^2^) of voltage-gated channels used in the model neuron. Ion channel densities were adjusted to reproduce observed experimental data. PNJ, paranodal junction; JXP, juxtaparanode; IND, internode; AIS, axon initial segment.

#### Stimulation

The ICLAMP point process in NEURON was used to inject current at different points in the neuron model. Based on Foutz et al.^79^, the photostimulus was modeled in Python and interfaced with the ChR2 model in NEURON. In brief, the ChR2 model has four states, two open (O1, O2) and two closed (C1, C2), whose rate of change is determined by

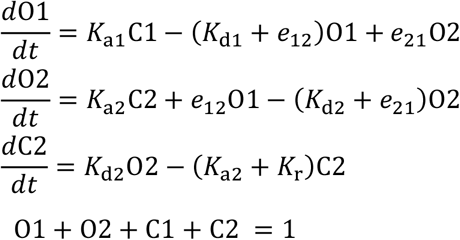

where *K*_d1_, *K*_d2_, *K*_r_, e_12_, and *e*_21_ are constants, whereas *K*_a1_ and *K*_a2_ depend on the flux of photons per unit area (Φ) according to

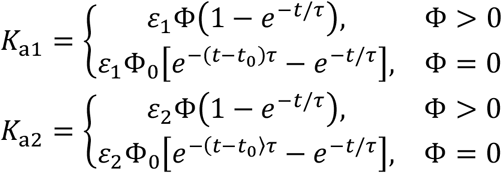

where *ε*_1_ and *ε*_2_ are quantum efficiency constants and τis a time constant for ChR2. During photostimulation, Φ is calculated as

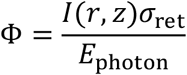

Where *σ*_ret_ is a channel cross section, *E*_photon_ is the energy of a photon, and *I*(*r, z*) is the irradiance *I* at each combination of radial distance *r* and depth *z*, as determined by

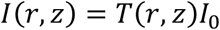

where *I*_0_ is irradiance of the light source and *T*(*r, z*) is the transmittance of light defined by the following function

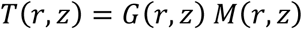

where *G*(*r, z*) is the Gaussian distribution of light, which in the absence of scattering is defined by

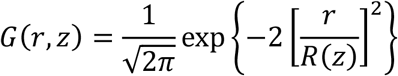

and *M*(*r, z*) depends on the properties of the tissue

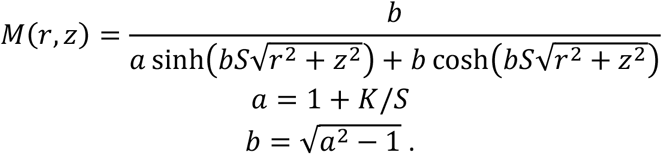

*S* (=0.125 mm^-1^) and *K* (=7.37 mm^-1^) are the scatter and absorption coefficients, respectively. *R*(*z*) is the radius of the column of light and was set to 10 to produce a photostimulation spot with diameter ∼20 μm. To model scattered light, a second term was added to *G*(*r, z*) to produce a distribution similar to Favre-Bulle et al.^80^.

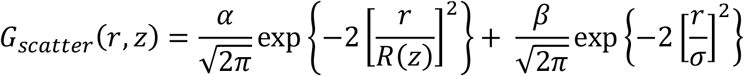

where α = 0.5, *β*= 0.00625 and *σ*= 800 to reproduce ChR2 activation data in Figure 1E. Light distributions are illustrated in **Figure S8**.

### Spike-triggered average and covariance analysis

In simulations, noisy current was injected into a node of Ranvier at the middle of the intact axon or at the middle of the AIS unless otherwise indicated. Noise was modeled as an Ornstein-Uhlenbeck process^81^,

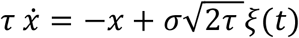

where *ξ*(*t*) was drawn from a Gaussian distribution with unit variance and mean *µ* = 0.1 nA, unless otherwise indicated. Noise was scaled by *σ* = 0.22 nA. The autocorrelation time *τ* is reported on the figures.

The original unfiltered noise was used for all analyses. The spike-triggered average (STA) was calculated as the average of all stimulus segments preceding a spike. The mean (*µ*) was subtracted from the STA to show stimulus fluctuations around zero. For the spike-triggered covariance (STC), the covariance matrix of the spike-triggered stimuli was formed based on the following equation^82^

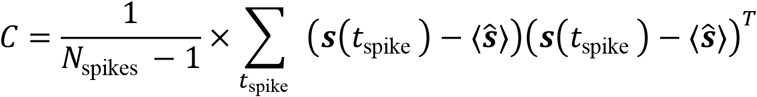

where 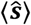 is the STA and *s*(*t*_spike_) is the stimulus segment preceding each spike. The covariance matrix of the surrogate data (*C*^*prior*^) was subtracted from *C* to find the covariance differences^83^

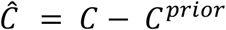

Surrogate data were obtained by shuffling the spike train^84^. Diagonalization of *Ĉ* reveals the significant eigenvalues and the corresponding eigenvectors (i.e. features). Significant eigenvalues were determined by comparing the eigenvalues of *C*^*prior*^ with the eigenvalues of *Ĉ*. The spike-triggered stimulus segments projected onto the features formed a distinct cloud that can be compared to the surrogate data projected onto the same features. At least 10,000 spikes were used for analysis.

### Axial current calculation

To calculate the axial current waveform a node receives during spike propagation, current was injected into the soma and the intracellular and extracellular voltages were recorded from a paranode adjacent to (just upstream of) a node in the middle of the axon (see cartoon on Fig. 4D). Axial current (*I*_1_) was measured in mA based on Ohm’s law

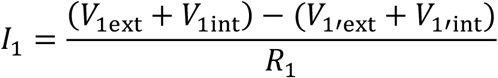

Results were validated using Kirchhoff’s current law,

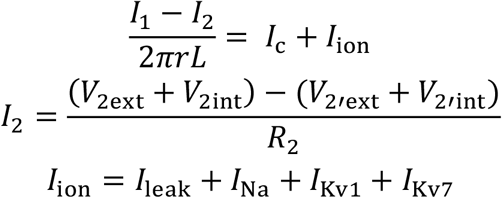

where *r* and *L* are the radius and length of the node, respectively. Any difference between the axial current flowing into and out of the node should be accounted for by capacitive and ionic current in the node.

## ACKNOWLEDGMENTS

This work was funded by a Canadian Institutes of Health Research Foundation Grant to SAP. MAK was supported by an Ontario Trillium Scholarship, Milligan Graduate Fellowship, Loo Geok Eng Foundation Scholarship, and Vanier Canada Graduate Scholarship.

## AUTHOR CONTRIBUTIONS

SAP conceived the study. MAK conducted experiments with assistance from SR. NA conducted simulations. All authors contributed to data analysis and prepared the manuscript.

## DECLARATION OF INTERESTS

The authors declare no conflicts of interest.

---

**Figure S1.**
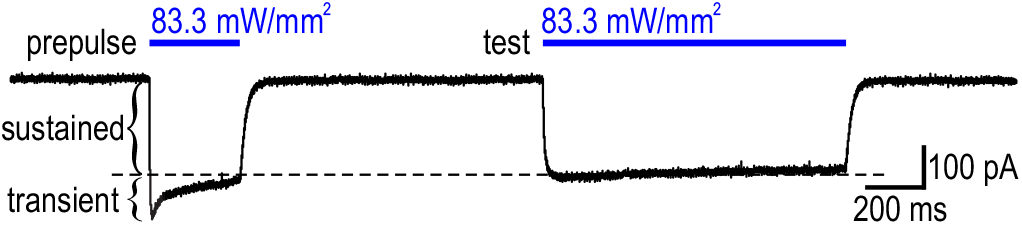
Removal of the transient (inactivating) component of ChR2 current. A photostimulus prepulse was applied before test photostimuli to remove the inactivating component of ChR2 current. Because ChR2 de-inactivates slowly, the subsequent (test) photostimulus evokes a “square” depolarizing current^40^. The prepulse is not depicted in other figures.

**Figure S2.**
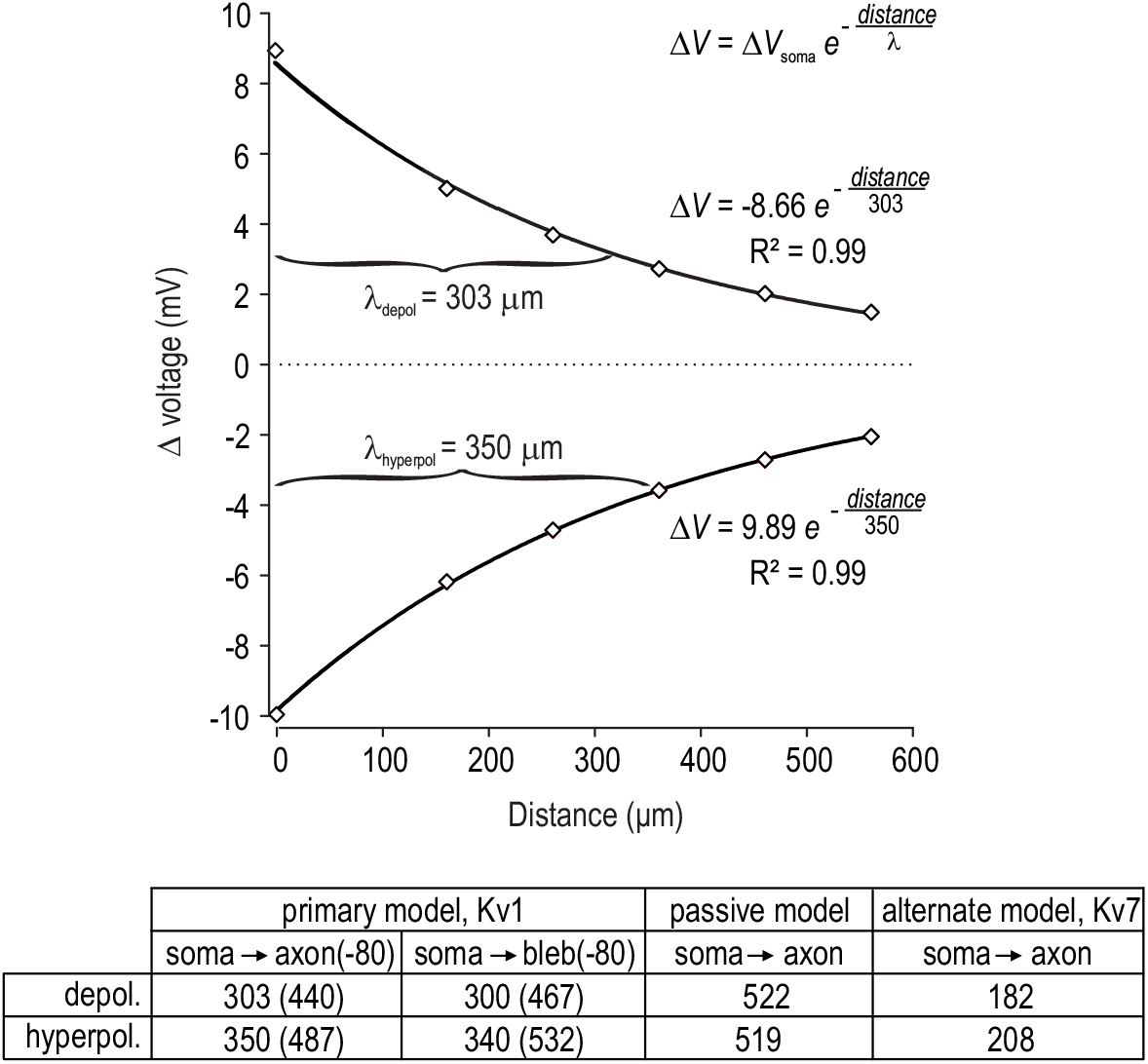
Axon length constant (λ) is shorter than previously reported. Estimates of the axonal length constant (λ) have typically ranged from 400 to 550 µm (refs. 53-55) but have reached as high as 800 µm (ref. 47). Some recent estimates are lower, in the range of 150 to 350 µm (refs. 91, 92). Using our model neuron, the voltage change (Λι*V*) caused by injecting depolarizing or hyperpolarizing current at the soma was measured at the soma and at several nodes located at different distances along the axon. Data were fit by the equation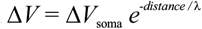, as illustrated in the top panel. The table summarizes λ (in μm) for different model conditions. Depolarizing and hyperpolarizing current yield similar values for λ in the passive axon, whereas depolarizing current spreads less than hyperpolarizing currents in active axons. The asymmetry in active axons is due to potassium currents activated by depolarization, which attenuate the spread of current; even hyperpolarizing current spreads less than in the passive axon because some potassium current is active at resting membrane potential^71^. Past models (*e*.*g*. ref. 54 based on ref.93) did not include low-threshold K_V_ 1 or K_V_ 7 channels. Experimental conditions are also a factor; for example, Alle and Geiger^53^ measured λλwhile hyperpolarizing the cell to -80 mV, which inactivates the aforementioned potassium channel and lengthens λ, according to our simulations. All values are based on measurements from resting membrane potential (−70 mV) except those indicated in brackets, where starting membrane potential was reset to -80 mV throughout the model. Experimental measurements using bleb recordings may overestimate λ because a sealed end forms where the axon is cut, which might restrict the flow of axial current. To explore this, we simulated bleb recording conditions by terminating the model axon at different lengths from the soma and measuring Λι*V* at the terminal, for each termination length. Values of thus obtained were not altered from values measured in the equivalent intact axon at -70 mV, and there was only a slight increase for measurements from -80 mV. Notably, voltage change (depolarization or hyperpolarization) attenuates more or less gradually depending on whether the underlying current is spreading toward or away from the soma (referred to in ref. 94 as *V*_in_ and *V*_out_, respectively). This means that λ measurements based on *V*_out_ (as described above) are longer than those based on *V*_in_ (ref. 21). To explain somatic depolarization based on axonal or dendritic ChR2 activation, λ measurements based on *V*_in_ are more appropriate, but electrotonic distance changes steeply near the junction between small neurites and the large soma because of the impedance mismatch^94^, rendering values based on *V* _in_less informative. For example, recording at the soma while injecting current in the axon 110 μm away, Shu et al.^18^ observed voltage deflections in the soma that were ∼10% as large as in the axon, indicating that current spread from the axon does not mediate strong somatic depolarization. By comparison, current arriving at the soma via multiple dendrites experience less impedance mismatch.

**Figure S3.**
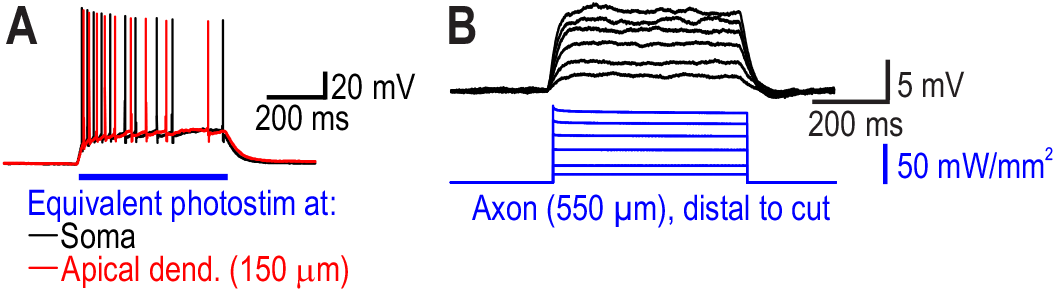
Experimental evidence for off-target ChR2 activation due to light scattering. **A**. Responses to photostimulation of equal intensity targeted to the soma (*black*) or apical dendrites (*red*) are similar. If the photostimulation zone were strictly focalized, the response (or photothreshold) should be sensitive to the precise positioning of the photostimulus onto the cell, but this was not observed. See Figure S4 for the effects of light scatter on the photothreshold in simulations. **B**. Photostimulation distal to a cut in the alveus still caused somatic depolarization proportional to the photostimulus intensity. This cannot be due to electrotonic spread of current from ChR2 activated in the photostimulation site since the axon is severed. Evidence that the space constant is shorter than previously reported, especially for the spread of depolarizing current (see Fig. S2), also argues against ChR2 current activated at the axon photostimulation site reaching the soma

**Figure S4.**
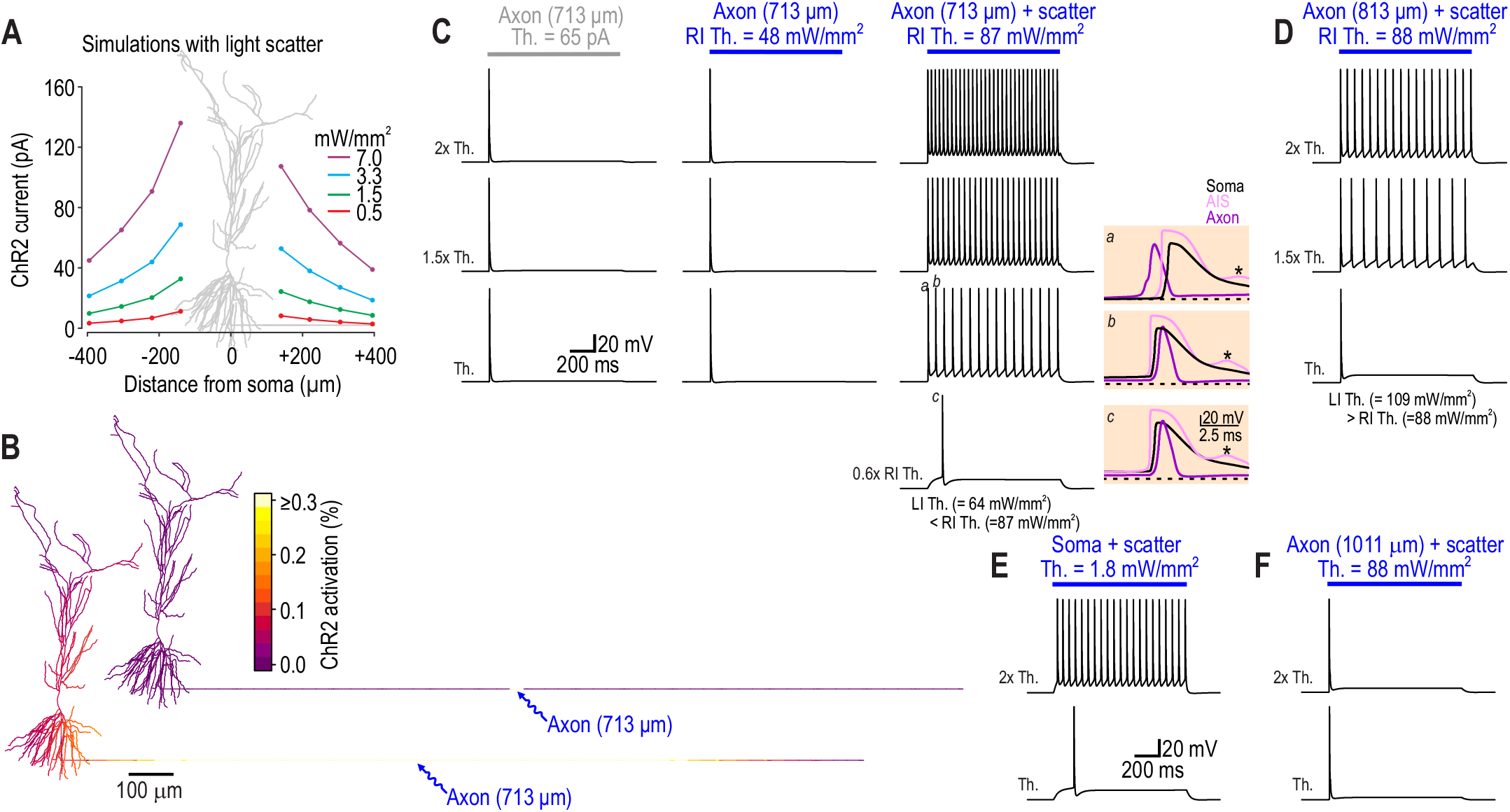
Impact of light scattering in simulations. **A**. ChR2 currents evoked by photostimulation at different distances from the soma toward (+) or away (−) from subiculum in the model neuron. Light scattering was tuned to reproduce experimental data in Figure 1E (see Methods. Neuron morphology is superimposed roughly to scale with the *x*-axis. In our model neuron, photostimuli in the – direction evoked slightly larger responses that photostimuli in the + direction; this was due to the dendritic geometry, as verified by simulations using a different morphology (data not shown). **B**. Color shows spatial distribution of ChR2 activated by axon photostimulation at 713 µm without light scattering (*top*) and with light scattering (*bottom*). In the latter case, ChR2 is activated in the dendrites and over a larger area of the axon, but whereas ChR2 activation reached 60% in the target zone, dendritic ChR2 activation was <0.3%. Color scale focuses on bottom range. Modest activation of dendritic ChR2 had a strong effect for multiple reasons. First, dendrites present a bigger target than the axon. Dendrites constitute 81% of the model neuron’s total surface area – compared with soma (2%) and axon (17%) – and branch extensively near the soma, unlike the axon which does not branch and whose surface area is thus distributed over a longer distance. CA1 pyramidal cell axons produce few collaterals near the soma^21^. Second, the soma and many dendrites are near the surface of the slice (since we used visually guided patching) whereas the axon may extend deep into the slice. Less light reaches deep structures because of scattering, especially in myelin-rich tissue like the alveus. These factors help explain why LI photothreshold (based on somatic photostimulation) is so much lower than RI photothreshold (based on axon photostimulation). **C**. Axon (713 µm) current injection (*left*) and photostimulation without light scattering (*center*) evoked only one RI spike and negligible somatic depolarization, whereas photostimulation with light scattering (*right*) evoked additional LI spikes and strong depolarization. Scattering caused RI photothreshold to increase from 48 to 87 mW/mm^2^. Photostimulation as weak as 64 mW/mm^2^ could evoke LI spikes (bottom trace). Beige insets show voltage at the soma (*black*), AIS (*pink*), and node adjacent to axon photostimulation site (*purple*). At RI photothreshold, the first spike (*a*) initiates in the axon whereas the next spike (*b*) and all subsequent spikes initiate in the AIS (*Beige insets*). At LI photothreshold, the spike (*c*) initiates in the AIS. * highlights failed spike initiation at the AIS (see Fig. S6A for example of bursts after blocking K_V_7 channels). **D**. Photostimulation farther down the axon (813 µm) was associated with the same RI photothreshold but a higher LI photothreshold than for photostimulation at 713 µm. Consequently, no LI spikes were evoked at RI photothreshold, unlike in C (compare also with experimental data in Fig. 2B). **E**. Light scatter did not alter the spiking pattern evoked by somatic photostimulation, but it increased the number of evoked spikes and reduced LI photothreshold (compare with left panel in Fig. 1C). The reduction in LI photothreshold contrasts the increase in RI photothreshold caused by scattering. **F**. When photostimulation with light scatter was shifted even farther down the axon (1011 µm), LI spikes were absent and somatic depolarization was attenuated but RI photothreshold was unchanged. The higher RI photothreshold observed with scattering is closer to RI photothrehsold values observed experimentally (see Fig. 2C). In the absence of scattering, RI photothreshold was sensitive to the exact positioning of the photostimulus along the model axon (data not shown), which is inconsistent with the subtler variability seen experimentally: RI photothreshold varied no more than 56% when re-tested at positions as far apart as 190 µm (the average RI photothreshold is reported in Fig. 2).

**Figure S5.**
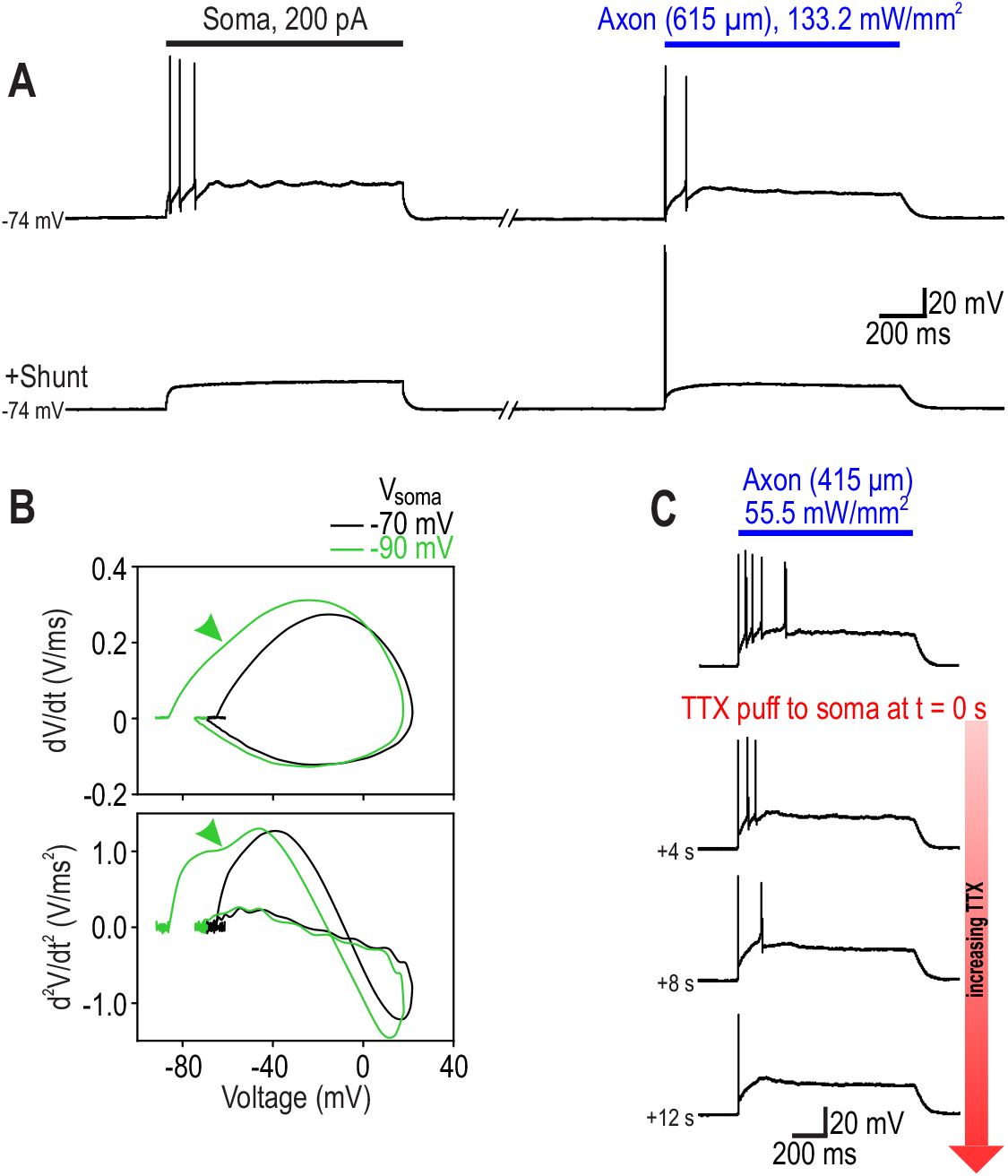
LI and RI spikes are differentially modulated by manipulations at the soma. **A**. A leak conductance (shunt) applied to the soma using dynamic clamp (*g*_leak_ = 5 nS, *E*_leak_ = -80 mV) eliminated LI spikes evoked by somatic current injection (*left*) and axon photostimulation (*right*). Only the RI spike evoked by axon photostimulation persisted. Input resistance was reduced by 35%, from 128 MΩ to 83 MΩ. **B**. Somatic hyperpolarization delayed activation of perisomatic sodium channels during somatic invasion of RI spikes. Phase portraits show voltage plotted against its velocity (d*V*/d*t*, top) and acceleration (d^2^*V*/d*t*^2^, bottom) during a typical RI spike. The spike recorded with *V*_soma_ = -90 mV (*green*) had a more pronounced inflection in its rising phase (*arrowhead*) than the spike recorded with *V*_soma_ = -70 mV (*black*). This inflection reflects the increased time required to charge the soma before activating perisomatic sodium channels when the starting voltage is more negative, and defines the IS-SD (initial segment-somatodendritic) latency^41, 67^. In our experiments, LI spikes had little if any IS-SD notch, which is consistent with past reports on CA1 pyramidal neurons^42^, but is unlike layer 5 pyramidal neurons^44^, where this notch is more prominent. **C**. Puffing TTX on the soma eliminated LI spikes more readily than RI spikes (see Fig. 2D). In 2 neurons, the single RI spike evoked by axon photostimulation persisted during somatic application of TTX, as illustrated here.

**Fig S6.**
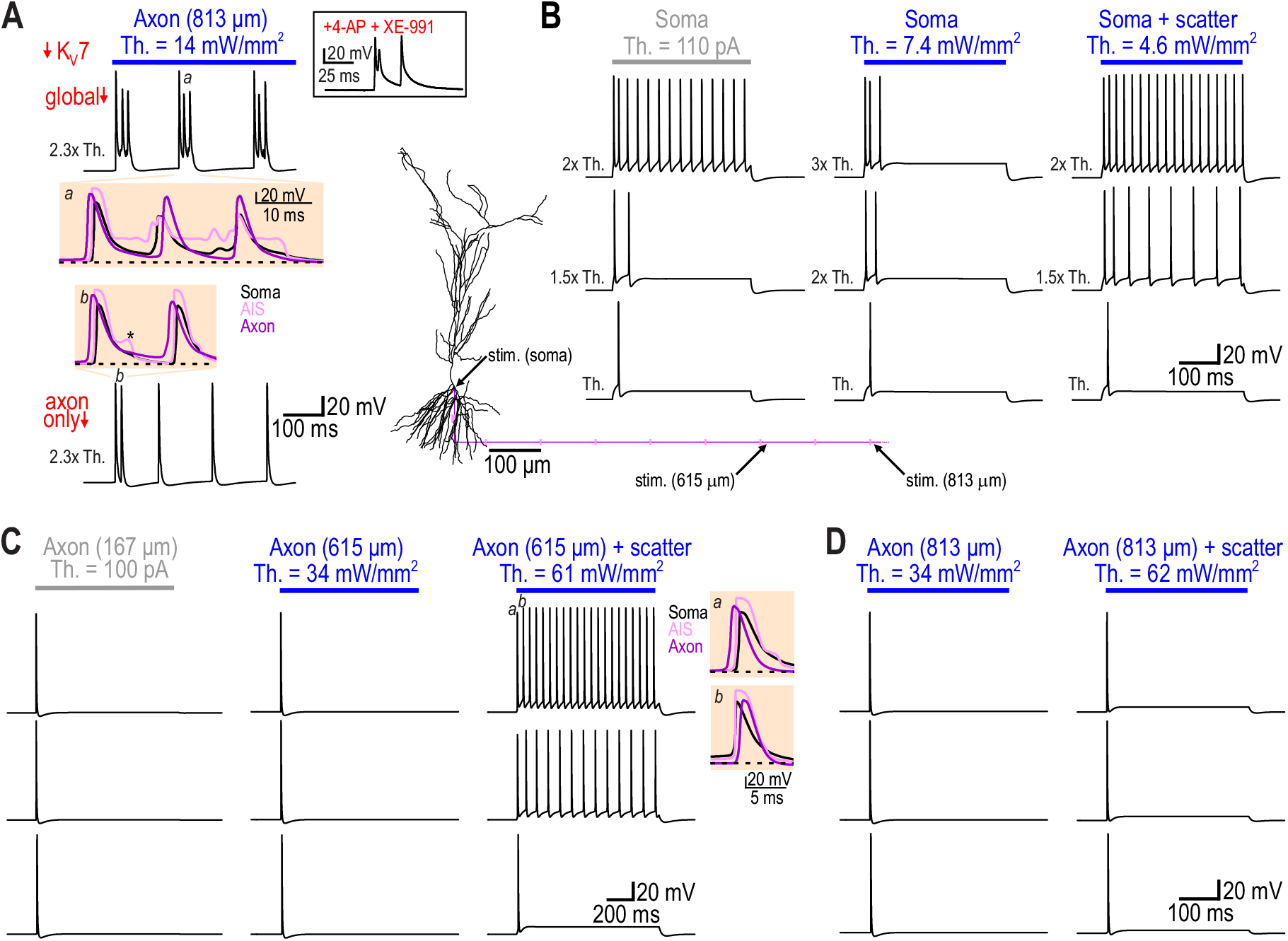
Similar axon response properties can be achieved with K_V_7 instead of K_V_1 channels. **A**. Reducing K_V_7 by 90% throughout an alternate model cell (*top*) allowed axon photostimulation to evoke repetitive RI spikes, but each RI spike was associated with a burst. Zoomed in view of the second burst (*beige inset*) shows an initial RI spike followed by two LI spikes resulting from sustained depolarization in the AIS. Similar bursts were occasionally observed experimentally in the presence of 30 µM XE-991 to block K_V_7 (box). Reducing K_V_7 selectively in the axon (bottom) caused repetitive RI spikes without bursting. Bottom beige inset shows that the initial pair of spikes both originate in the axon. The bump on the falling phase of AIS spikes (*) is still evident when K_V_7 is selectively reduced in the axon (see also Fig. S4C), but it is attenuated compared with global reduction of K_V_7. These results argue that K_V_7 regulates perisomatic excitability (see bottom trace of Fig. 3D). It is, however, notable that K_V_7 could serve the same role as K_V_1 in regulating axonal excitability, as demonstrated in B-D. **B**. When applied at the soma, current injection (*left*) and photostimulation (*center, right*) all evoked repetitive LI spiking. Light scatter had the same effect as in our original model (see Fig. S4), namely increasing the evoked spiking and reducing LI photothreshold. **C**. When applied at the axon (615 µm), current injection (*left*) and photostimulation without light scattering (*center*) evoked a single RI spike, but additional LI spikes were produced when light scattering was included (*right*). Beige insets confirm that the first spike (*a*) originated in the axon whereas the second spike (*b*) originated in the AIS. Like in the original model (see Fig. S4), light scatter increased RI photothreshold. In this particular condition, LI photothreshold was slightly higher than RI photothreshold. **D**. When applied farther down the axon (813 µm), photostimulation with and without light scatter (*right* and *center*, respectively) both evoked a single RI spike.

**Fig S7.**
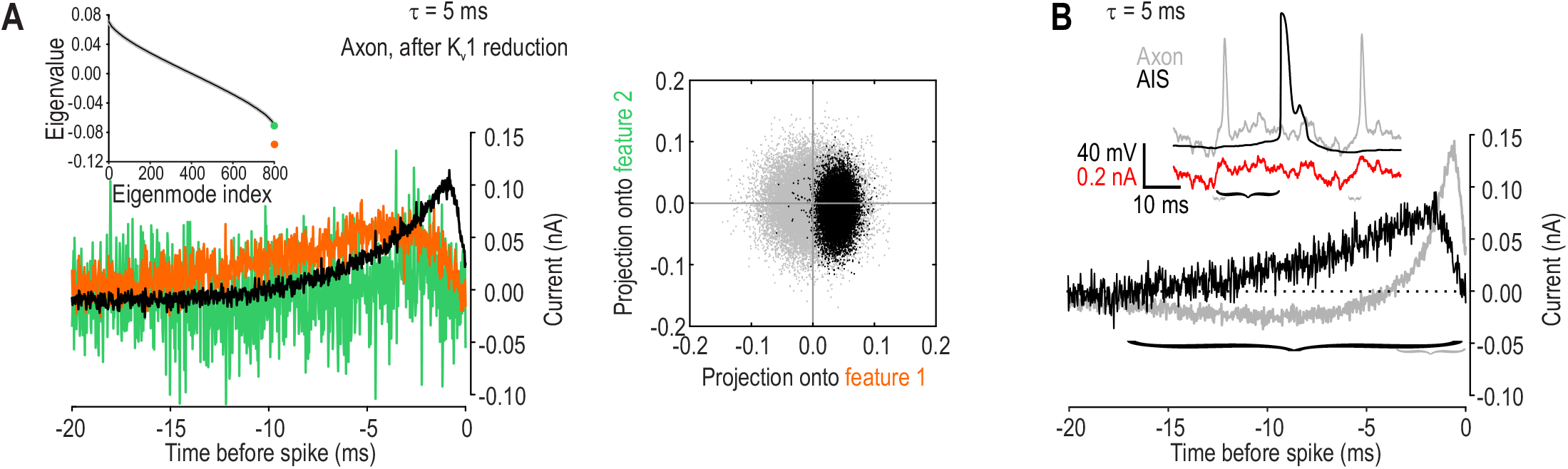
Filtering in the axon and AIS. **A**. K_V_1 channels are necessary for the high-pass filtering exhibited by the axon. Analysis like in Figure 4C was repeated but with K_V_1 density reduced by 90%. There is only one significant eigenvalue and the broad monophasic STA resembles results from the AIS more closely than it does the narrow biphasic STA of the normal axon. **B**. The axon and AIS respond to different features of the noisy input. Traces (*top*) show sample voltage responses at the axon (*gray*) and AIS (*black*) to noisy current (*red*, τ = 5 ms, χρ = 0.22 nA and µ = 0.1 nA) injected at the axon or soma, respectively. Noise was injected at the soma rather than directly into the AIS to model the effect of synaptic current passing through the soma to reach the AIS, and being low-pass filtered on the way. Note that the voltage response in the AIS is much smoother than in the axon. Whereas abrupt increases in input (*gray brackets*) drive the axon to spike, a sustained increase (*black bracket*) is required to drive the AIS to spike. Brackets are shown to scale underneath STAs (*bottom*) calculated from at least 10,000 spikes for the corresponding condition. This noise does not represent the input typically experienced by the axon but it is nonetheless useful for characterizing the axon response; Ornstein-Uhlenbeck noise generated with τ = 5 ms is near the top end of frequencies the AIS/soma can respond to, but is at the bottom end of frequencies the axon can respond to (see Fig. 4F).

**Fig S8.**
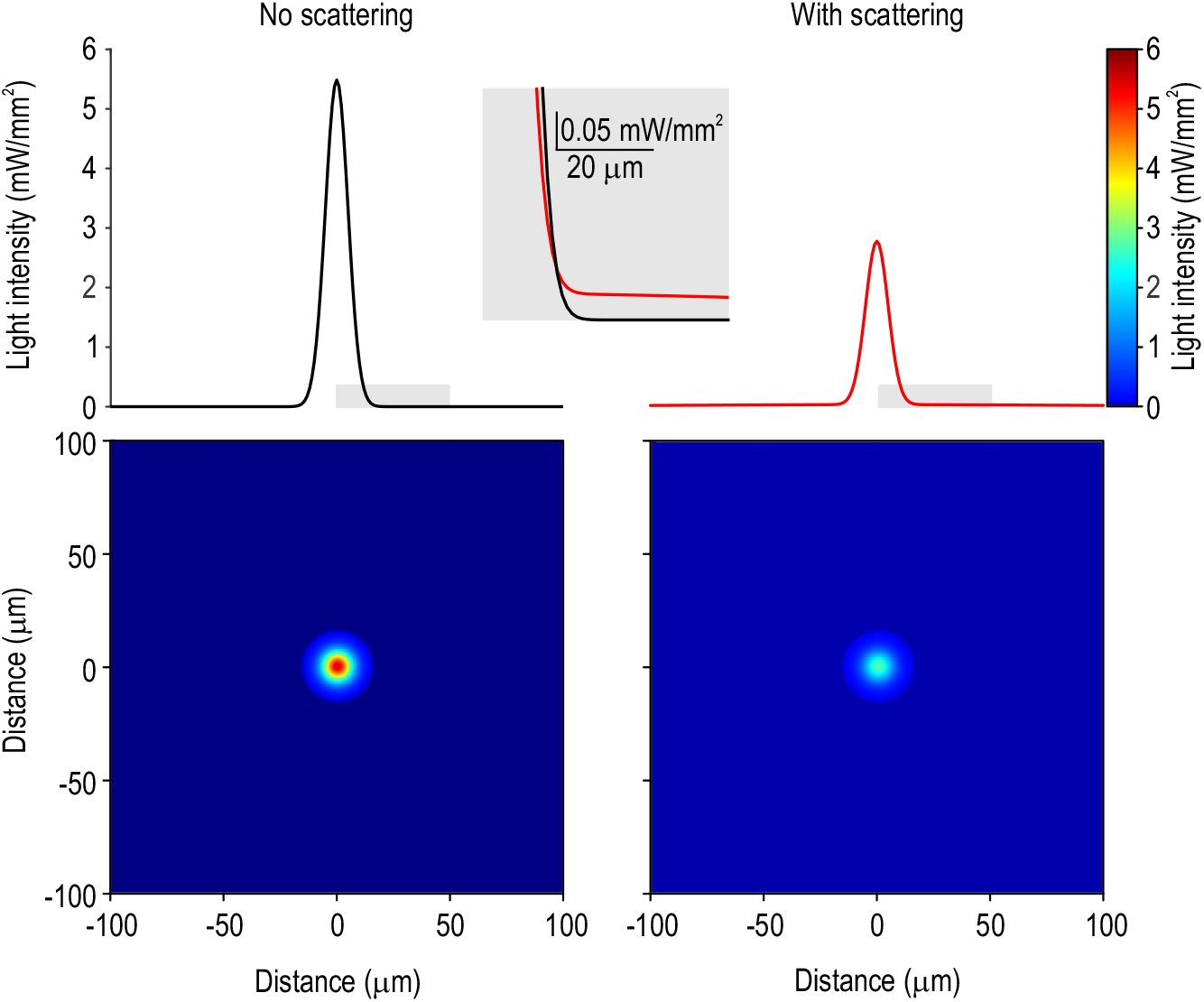
Distribution of photostimulation light in the computational model. **A**. Distribution of photostimulation light with (*right*) and without *(left*) scattering included in the light distribution model. The distribution without scatter was set based on the size of the blue light spot visible during photostimulation after partially closing the field diaphragm (see Methods). Starting from that distribution, a fraction of light was re-allocated to a broader distribution adjusted to reproduce voltage clamp data in Figure 1E (see Fig. S4A). The total amount of light (corresponding to total area under the curve) is equivalent between conditions before accounting for absorption. The scattered light is represented by the long tails of the red curve; gray inset shows zoomed in view of gray regions highlighted on main graphs. The scattered light is evident as a very subtle variation in blue on the 2D distributions (*bottom*). Intensity is controlled by scaling distributions by *I*_0_ (= 20 mW/mm^2^ for examples shown). Intensity is reported as the integral over space.

**Table S1.** Na^+^ channel reduction (% from normal) at time points i-v in Figure 2F.

| Time point | AIS | Axon hillock | Soma | Dendrites |  |  |
| --- | --- | --- | --- | --- | --- | --- |
|  |  |  |  | Primary, including apical <50 µm from soma | Secondary <65 µm from soma and apical >50 µm from soma | Secondary & tertiary >65 µm from soma |
| i | 10 | 20 | 30 | 20 | 10 | 0 |
| ii | 30 | 40 | 50 | 40 | 30 | 20 |
| iii | 50 | 60 | 70 | 60 | 50 | 40 |
| iv | 60 | 70 | 80 | 70 | 60 | 50 |
| v | 70 | 80 | 90 | 80 | 70 | 60 |

## Notes

### Competing Interest Statement

The authors have declared no competing interest.

